# The Amplified Burden of Depression in Drug-Resistant Epilepsy: Role of Limbic Network Dynamic Reconfiguration

**DOI:** 10.1101/2025.07.20.665759

**Authors:** Zhoukang Wu, Haiyun Tang, Liangjiecheng Huang, Min Wang, Yaotian Gao, Aobo Chen, Hang Cao, YuanJin Gong, Jia Hu, Xiaoping Yi, Xiaosong He

**Affiliations:** Department of Psychology, State Key Laboratory of Eye Health, University of Science and Technology of China,Hefei 230026, China; Department of Radiology, Xiangya Hospital, Central South University, Changsha, Hunan, 410078, China; Department of Neurosurgery, The First Affiliated Hospital of USTC, Division of Life Sciences and Medicine, University of Science and Technology of China, Hefei, Anhui, 230001, China; Department of Neurosurgery, Xuanwu Hospital, Capital Medical University, Beijing, 100053, China; Department of Radiology, Chongqing University Three Gorges Hospital, Chongqing University, Chongqing 410008, Chongqing, P.R. China; School of Medicine, Chongqing University, Chongqing 400030, P.R. China; Clinical Research Center (CRC), Medical Pathology Center (MPC), Cancer Early Detection and Treatment Center (CEDTC) and Translational Medicine Research Center (TMRC), Chongqing University Three Gorges Hospital, Chongqing University, Chongqing 404000, P.R. China; National Clinical Research Center for Geriatric Disorders (Xiangya Hospital), Central South University, Changsha 410008, Hunan, P.R. China; FuRong Laboratory, Changsha 410078, Hunan, P.R. China

## Abstract

**Objectives:** To explore potential associations between drug resistance and limbic network (LN) dynamic functional interactions in temporal lobe epilepsy (TLE), whether these LN alterations are associated with and potentially mediate comorbid depression severity, and their potential neurochemical and transcriptomic underpinnings.

**Methods:** This cross-sectional study included 49 patients in the Responsive group, 33 patients in the Resistant group, and 50 healthy controls (HCs). Using resting-state fMRI, we derived LN dynamic integration coefficients and used non-negative matrix factorization (NMF) to identify coactivation patterns. Depressive symptoms (Self-Rating Depression Scale, SDS), LN integration, and NMF pattern expression were compared across groups (ANOVA) and correlated. Mediation analysis tested if LN integration mediated the relationship between drug resistance status and SDS scores. LN integration differences were spatially correlated with neurotransmitter receptor density and Allen Human Brain Atlas transcriptomic data.

**Results:** Patients in the Resistant group reported significantly higher SDS scores than patients in the Responsive group (*p* = 0.008). LN integration with the dorsal attention network (DAN) and fronto-parietal network (FPN) was significantly lower in the Resistant group compared to the Responsive group (*p* = 0.001; *p* < 0.001). NMF analysis identified key LN-DAN (*p* = 0.013) and LN-FPN (*p* < 0.001) coactivation patterns. In TLE patients, DAN-LN integration was negatively correlated with depressive symptoms (*r* = -0.28, *p* = 0.01), and lower integration levels significantly mediated the relationship between group status (Resistant vs. Responsive) and increased symptom severity (*p* < 0.001). These LN integration coefficient differences (Responsive group vs. Resistant group) were associated with 5-hydroxytryptamine 1B (5-HT1B; *r* = 0.29, *p* = 0.036) and dopamine D2 (D2; *r* = -0.29, *p* = 0.034) receptor densities, and linked to gene expression pathways including telomerase activity regulation (*p* < 0.001).

**Interpretation:** This study identifies dynamic LN dysregulation, supported by distinct neurochemical and transcriptomic profiles, as a core bridging mechanism potentially contributing to comorbid depression linked to drug response status in TLE patients.

## Introduction

Depression is the most common psychiatric comorbidity in epilepsy, with a prevalence ranging from 20% to 60% among patients with drug-resistant epilepsy (DRE), such as those with temporal lobe epilepsy (TLE).^1, 2^ This comorbidity significantly impairs seizure control, reduces adherence to antiseizure medications, and diminishes quality of life.^3, 4^ The relationship between epilepsy and depression is considered bidirectional,^5, 6^ potentially stemming from shared pathogenic mechanisms,^7^ with the limbic system being a key area implicated in both conditions due to its role in emotional regulation. ^8, 9^ Indeed, structural changes in the amygdala and hippocampus,^10,11^ as well as altered functional connectivity of the limbic system, particularly involving the hippocampus and its connections to emotional-related brain regions,^12^ have been reported in epilepsy patients with depression.

Despite evidence implicating limbic system alterations in the comorbidity of epilepsy and depression, particularly in DRE, it remains unclear how drug resistance itself specifically modifies or exacerbates depressive symptoms through these neural substrates. While psychiatric disorders, including depression, are associated with drug resistance,^13–15^ the neurobiological mechanisms—such as neurotransmitter imbalances or specific functional brain changes—by which drug resistance might worsen depression are not well understood. Current hypotheses on drug resistance often focus on biological factors like altered drug metabolism or genetic variations and neural network abnormalities,^16, 17^ but largely overlook the implications for psychiatric comorbidities.^18^ Consequently, there is a paucity of neuroimaging studies investigating how drug resistance and depressive symptoms are linked in DRE through specific neural biomarkers, with most research examining either epilepsy or psychiatric conditions in isolation.

Dynamic functional connectivity (dFC) has emerged as a valuable approach for studying brain state fluctuations and has revealed significant dFC abnormalities in depression, such as altered connectivity variability^19, 20^ and inter-network integration.^21^ Given that epilepsy is also characterized by dynamic neural activity, including excessive synchrony,^22^ dFC analysis offers a novel perspective to investigate how drug resistance in DRE might be associated with an exacerbation of comorbid depressive symptoms through potential disruptions in these brain dynamics.

The purpose of the present study was to utilize dFC analysis of resting-state fMRI data, integrated with transcriptomic and neurotransmitter density information, to explore potential associations between drug response status in TLE and the dynamic functional interactions of the limbic network (LN). We aimed to elucidate whether altered dynamic LN reconfigurations are associated with and potentially mediate the relationship between this group status (Resistant vs. Responsive) and the severity of comorbid depressive symptoms, and to explore the neurochemical and genetic underpinnings potentially linked to these processes.

## Methods

### Participants

This cross-sectional study utilized fMRI data collected at Xiangya Hospital, Central South University, between January 1, 2019, and December 21, 2021. The study protocol was approved by the Ethics Committee of Xiangya Hospital, Central South University (Approval No. 201803420), and adhered to the Declaration of Helsinki; all participants provided written informed consent.

Patients with TLE were screened and recruited by two experienced radiologists and two neurologists from the neurology outpatient clinic. Inclusion criteria for TLE patients were: (1) long-term video-electroencephalography (vEEG) indicating temporal lobe origin of seizures; (2) seizure semiology consistent with mesial temporal lobe epilepsy (mTLE); and (3) MRI findings negative for pathologies other than hippocampal sclerosis (HS). Patients with TLE were further categorized based on their response to antiseizure medication (ASM), which enabled comparative analyses of depressive symptoms and associated neural mechanisms across different treatment response profiles.

HCs were recruited from the local community and had no history of neurological or psychiatric disorders, nor contraindications for MRI. Clinical depressive symptoms in Participants were assessed using the Self-Rating Depression Scale (SDS).^23^

### Definitions

Drug-resistant epilepsy (DRE) was defined according to the 2010 International League Against Epilepsy (ILAE) criteria^24^ as the failure of adequate trials of two tolerated, appropriately chosen and used antiseizure medication (ASM) schedules (monotherapies or in combination) to achieve sustained seizure freedom. The Resistant group consisted of TLE patients meeting these ILAE criteria for DRE at enrollment. The Responsive group included TLE patients who were seizure-free for at least one year on a stable ASM regimen at enrollment and did not meet ILAE criteria for DRE.

### Imaging Acquisition and Preprocessing

Resting-state fMRI data were acquired on a Siemens 3.0 T Prisma MRI system at Xiangya Hospital. Participants were instructed to remain awake with eyes closed and relaxed during scanning; none were reported to have fallen asleep. Scan parameters were: repetition time (TR) = 2.00 s, echo time (TE) = 30 ms, flip angle = 66°, 75 slices, slice thickness = 2 mm (no gap), field of view = 220 × 220 mm², voxel size = 2.34 × 2.34 × 2 mm³, matrix = 94 × 94, 239 volumes, and a multiband acceleration factor of 3.

Functional images were preprocessed using fMRIprep,^25^ including estimation of head motion parameters, slice-timing correction, resampling to native space for motion correction, co-registration to T1-weighted anatomical images, and normalization to Montreal Neurological Institute (MNI) standard space. Post-processing was performed using XCP-D,^26^ involving framewise displacement (FD) calculation for motion spike identification and despiking, denoising with 24 nuisance regressors, bandpass filtering (0.01-0.1 Hz), censoring of timepoints with FD > 0.5 mm, and spatial smoothing (Gaussian kernel, 6mm FWHM). (See Fig. 1 for an overview of preprocessing and analysis steps).

**Figure 1.**
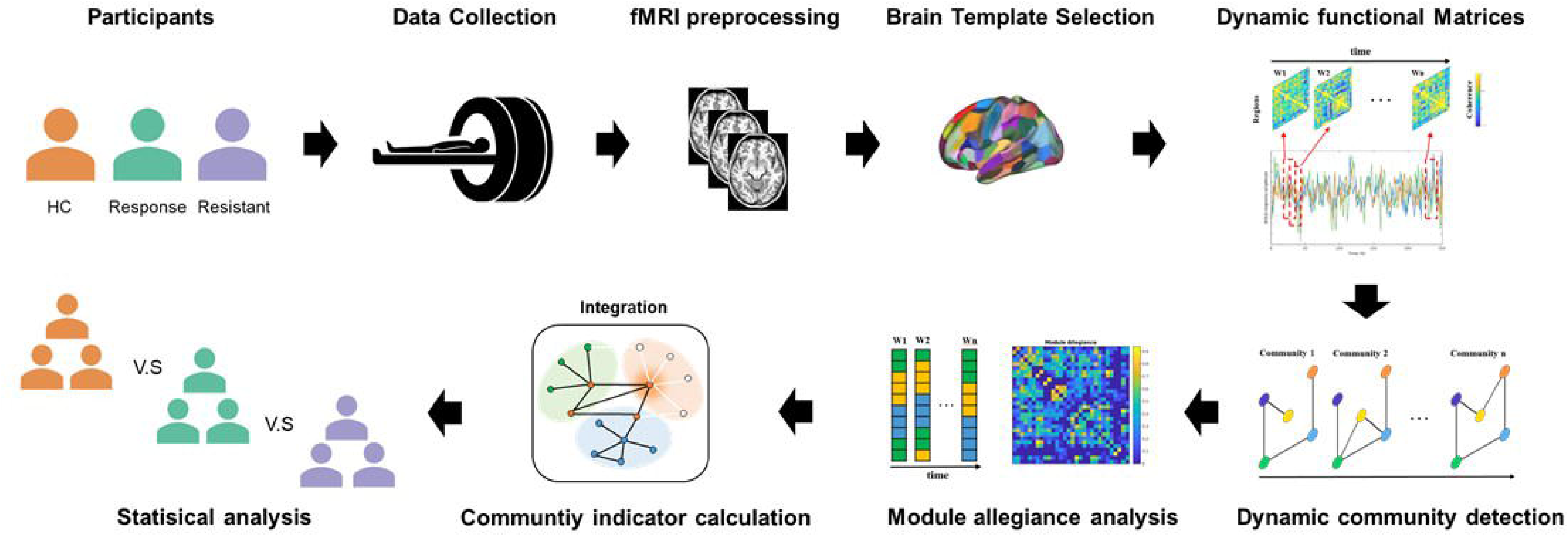
Core data analysis pipeline of the current study This figure illustrates the key analytical steps from fMRI data preprocessing to the extraction of individualized dynamic network features and subsequent statistical modeling. TLE patients and HCs were recruited, and the TLE patients were further subdivided into the Responsive group and the Resistant group based on their treatment outcomes. All participants underwent fMRI scanning. The acquired data were subjected to standardized preprocessing, followed by cortical parcellation and extraction of regional time series. Dynamic functional connectivity (dFC) matrices were then constructed using a sliding-window approach. Modularity-based algorithms (e.g., module allegiance matrices) were employed to extract network integration features across time windows. Finally, statistical modeling was used to systematically compare network characteristics across different groups.

### Statistical Analysis of Clinical Data

Clinical and demographic data were analyzed using SPSS Statistics 24.0 (IBM Corp., Armonk, NY). Continuous variables are presented as mean ± standard deviation (SD) or median (interquartile range), according to their data distribution. To compare demographic and clinical data between study groups, comparisons of continuous variables (e.g., SDS scores, age) between two independent groups were made using independent samples t-tests, while comparisons among the three groups (HC, Responsive group, Resistant group) were performed using one-way analysis of variance (ANOVA) followed by post-hoc two-sample t-tests. If assumptions for these parametric tests were not met, Mann-Whitney U tests (for two groups) or Kruskal-Wallis tests with appropriate post-hoc comparisons (for three or more groups) were employed. For categorical variables (e.g., gender), comparisons were made using Pearson’s χ² test or Fisher’s exact test, as appropriate. A significance level of *p* < 0.05 (two-tailed) was used for these initial group comparisons (univariate analyses).

### Dynamic Functional Connectivity Analysis

Dynamic network analysis was conducted using the Gretna toolbox.^27^ Brain networks were constructed using the BNA246 template (246 regions of interest, ROIs),^28^ with ROIs assigned to the Yeo 7-networks plus a subcortical network (SUB). BOLD time-series were divided into consecutive windows (length: 15 TRs, step: 1 TR). Dynamic functional connectivity matrices (246 × 246) were calculated for each window, with negative connections set to zero, and then linked to form a multilayer network.

Multilayer community detection was performed using the GenLouvain toolbox, employing a generalized Louvain algorithm with a multilayer modularity quality function^29^ (structural resolution *γ* = 1 and temporal resolution *ω* =1). To ensure stability, 100 iterations were performed per subject. ^30^ The dynamic metric "integration" was calculated to quantify interactions between networks, representing the average probability that a node is in the same community as nodes from other networks. Network-level integration was defined as the average integration coefficient of ROIs within that network.

To evaluate the impact of drug resistance on LN dynamic activity, one-way ANOVA was used to compare LN integration coefficients (LN-all, LN-DAN, LN-FPN) among the HC, Responsive group, and Resistant group. Post-hoc two-sample t-tests with Bonferroni correction (P < 0.05) were used for pairwise comparisons. The relationship between LN integration coefficients and SDS scores in TLE patients was explored using Pearson correlation analysis (P < 0.05). Subsequently, mediation analysis (PROCESS for SPSS v24.0)^31^ with bootstrapping (5000 samples) and bias-corrected 95% confidence intervals (CI) was employed to investigate if LN dynamic reconfiguration (specifically DAN-LN integration) mediated the relationship between drug resistance status (Responsive group vs. Resistant group) and depressive symptoms. A significant mediation effect was indicated if the 95% CI did not include zero.

### 2.6 Non-negative Matrix Factorization (NMF) Analysis

NMF was applied to ROI-level integration coefficient matrices of LN-DAN and LN-FPN to identify representative coactivation patterns (features) and their individual expression levels.^32^ The optimal number of components (k) was determined using the lcasso framework by evaluating the stability index (Iq) of the decomposition across multiple runs. These NMF-based processing steps were implemented using custom MATLAB scripts, which are based on the stability-driven NMF toolbox^33^ and the Collaborative Brain Decomposition and Brain Decoding Packages.^34^ Differences in the expression levels of NMF-identified coactivation patterns among the three groups were assessed using one-way ANOVA, followed by post-hoc two-sample t-tests with Bonferroni correction (P < 0.05).

### Spatial correlation analyses for neuroimaging and neurotransmitter density

To identify neurotransmitters associated with group differences in LN integration, spatial Pearson correlation analysis was conducted between t-statistic maps reflecting differences in LN integration coefficients (LN-DAN, LN-FPN; Responsive group vs. Resistant group) and 19 neurotransmitter receptor density maps.^35^ The statistical significance of these correlations was assessed using a spatial permutation test (1,000 rotations), and the resulting p-values were subsequently corrected for multiple comparisons using the False Discovery Rate (FDR) method. These receptor maps were accessed from a publicly available repository (https://github.com/netneurolab/hansen_receptors/tree/main/data/PET_nifti_images).

### Neuroimaging-transcriptome spatial association analysis

To explore the molecular basis of functional differences, neuroimaging-transcriptomic spatial association analysis was performed. Gene expression data from the Allen Human Brain Atlas (AHBA; http://www.brain-map.org; 6 postmortem donors)^36, 37^ were processed by selecting representative probes (expression intensity >50% of background; removing duplicates and probes with Spearman Rho < 0.2 to RNA-seq data),^38^ resulting in 15,633 genes. Gene expression data were projected onto the BNA246 brain atlas. The ABAnnotate toolbox^39^ was then used to assess the spatial association between these gene expression profiles and the t-statistic maps of LN integration differences (focused on 52 ROIs from DAN and FPN), and to identify potentially enriched gene categories and molecular pathways. The statistical significance of the enrichment for these identified categories and pathways was assessed using spatial permutation testing, which involved 1,000 rotations of the LN integration t-map to generate a null distribution, and p-values were subsequently FDR-corrected.

## Results

### Clinical characteristics

Initially, 50 TLE patients were recruited for the Responsive group and 35 for the Resistant group. One patient from the Responsive group was excluded for high signal dropout, and two from the Resistant group for high motion artifact. The final cohort included 49 patients in the Responsive group, 33 in the Resistant group, and 50 HCs.

Demographic and detailed clinical characteristics are presented in Table 1. The three groups were well-matched for age (F(2,129) = 0.016, *p* = 0.984) and gender (χ²(2) = 0.911, *p* = 0.634). Key epilepsy-related characteristics, including age at onset, duration of epilepsy, seizure focus distribution, and baseline seizure severity, did not significantly differ between the Responsive group and Resistant group (all *p* > 0.05; see Table 1).

**Table 1.** Demographic and clinical characteristics of the used subjects.

| Characteristics | HC(50) | Responsive group (49) | Resistant group (33) | $F/t/\chi^2$ value | HC vs. Responsive group | HC vs. Resistant group | Responsive group vs. Resistant group |
| --- | --- | --- | --- | --- | --- | --- | --- |
| Age(years) | 28.46±13.77 | 28.67±11.89 | 28.18±10.47 | $F_{129} = 0.016$<br>($p = 0.984$) | $p = 0.931$ | $p = 0.920$ | $p = 0.86$ |
| Gender, male/female | 25/25 | 23/26 | 19/14 | $\chi^2_2 = 0.911$<br>($p = 0.634$) | / | / | / |
| SDS scores | 33.68±4.24 | 47.16±9.52 | 51.76±8.40 | $F_{129} = 66.47$<br>($p < 0.001$) | $p < 0.001$ | $p < 0.001$ | $p = 0.008$ |
| Age at epilepsy onset | / | 21.11 ± 12.42 | 18.00 ± 10.52 | $t_{80}=1.18$ ($p = 0.241$) | / | / | / |
| Duration of epilepsy | / | 7.44 ± 7.12 | 10.04 ± 8.73 | $t_{80}=-1.48$ ($p = 0.143$) | / | / | / |
| Seizure focus, Left/Right/All | / | 29/17/3 | 17/13/3 | $\chi^2_2 = 0.563$ ($p = 0.755$) | / | / | / |
| Baseline seizure severity score | | 7.49 ± 1.66 | 6.94 ± 1.64 | $t_{80}=1.48$ ($p = 0.143$) | / | / | / |

Note: One-way analysis of variance (ANOVA) was first used to determine the differences in
demographic and clinical characteristics. Post-hoc two-sample t-tests were then used to identify group
differences in all clinical measures. Gender comparisons were conducted using Pearson's chi-square test.
Abbreviations: SDS, Self-Rating Depression Scale; HC, Healthy Control.

Self-Rating Depression Scale (SDS) scores differed significantly among groups (F(2,129) = 66.47, *p* < 0.001). Both the Responsive group (47.16 ± 9.52) and the Resistant group (51.76 ± 8.40) groups reported higher SDS scores than HCs (33.68 ± 4.24) (both *p*_Bonferroni_ < 0.001). Notably, the Resistant group exhibited significantly higher SDS scores than the Responsive group (*p*_Bonferroni_ = 0.008), indicating a greater burden of depressive symptoms in drug-resistant TLE patients.

### Network Integration differences among HC, Responsive group, and Resistant group

ANOVA analysis revealed significant differences in integration coefficients among the HC, Responsive group, and Resistant group in the LN-all (*F*_(2, 129)_ = 25.70, *p* < 0.001, see Fig.2A), LN-DAN (*F*_(2, 129)_ = 9.91, *p* < 0.001, see Fig.2B), and LN-FPN (*F*_(2, 129)_ = 10.06, *p* < 0.001, see Fig.2C). Post hoc two-sample t-tests further indicated that the integration coefficients in the Resistant group were significantly lower than those in the Responsive group after Bonferroni correction(LN-all: *t* = -4.14, *p*_Bonferroni_ < 0.001, see Fig.2A; LN-DAN: *t* = -3.67, *p*_Bonferroni_ = 0.001, see Fig.2B; LN-FPN: *t* =-3.88, *p*_Bonferroni_ < 0.001, see Fig.2C). Additionally, post hoc tests showed no significant differences in the integration coefficients of the LN-DAN (t = -0.597, *p*_Bonferroni_ = 1, see Fig2.B) and LN-FPN (*t* = -0.267, *p*_Bonferroni_ = 1, see Fig2.C) networks between the HC and Responsive group. The above results suggest that drug response status in TLE may be closely related to alterations in the dynamic integration capacity of the LN.

**Figure 2.**
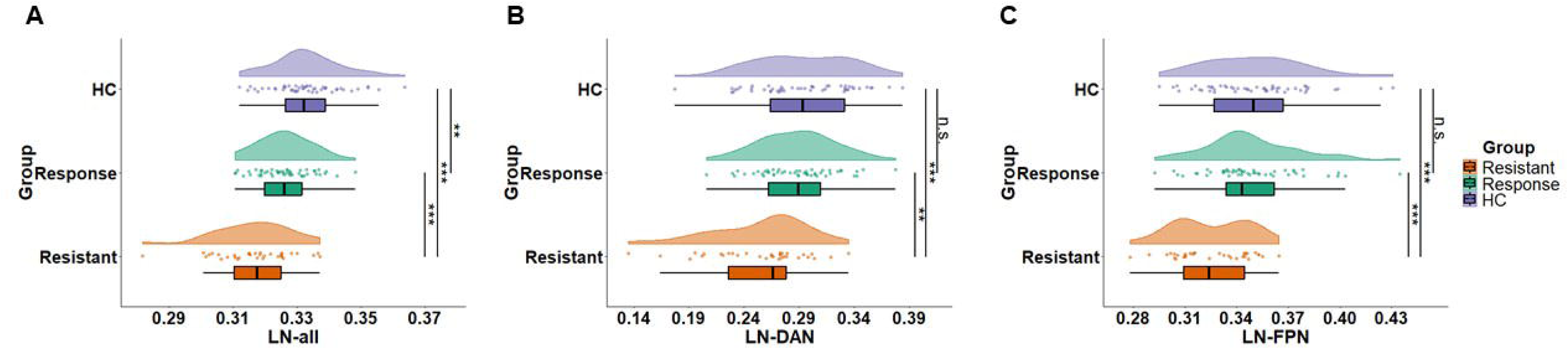
Group differences in integration coefficients in the LN (A) Integration between LN and other brain networks (LN–all): ANOVA revealed a significant group effect among HC, the Responsive group, and the Resistant group (*F*_(2, 129)_ = 25.70, *p* < 0.001). Post hoc two-sample t-tests showed significantly lower integration coefficients in the Resistant group compared to the Responsive group (*t* = −4.14, *p*_Bonferroni_ < 0.001). (B) Integration between LN and the dorsal attention network (LN–DAN): ANOVA revealed significant group differences (*F*_(2, 129)_ = 9.91, *p* < 0.001). The Resistant group showed significantly reduced integration compared to the Responsive group (*t* = −3.67, *p*_Bonferroni_ = 0.001). No significant difference was found between HC and the Responsive group (*t* = −0.60, *p*_Bonferroni_ = 1). (C) Integration between LN and the frontoparietal network (LN–FPN): ANOVA indicated a significant group effect (*F*_(2, 129)_ = 10.06, *p* < 0.001). Post hoc comparisons showed that the Resistant group had significantly lower integration than the Responsive group (*t* = −3.88, *p*_Bonferroni_ < 0.001). No significant difference was observed between HC and the Responsive group (*t* = −0.27, *p*_Bonferroni_ = 1). \**p* < 0.05, \*\**p* < 0.01, \*\*\**p* < 0.001.

### Identification of Classic Coactivation Patterns

To explore whether similar results exist at the ROI level as at the network level, we used NMF to identify 4 classic coactivation patterns in the ROI-level integration coefficient matrix of the LN-DAN. One-way ANOVA results showed that pattern 1 (pattern#1) exhibited significant differences between the Responsive group, Resistant group, and HC group (*F*_(2, 129)_ = 5.823, *p* = 0.004, see Fig.3B). Further post-hoc two-sample t-tests indicated that, after Bonferroni correction, the integration coefficient of the Resistant group was significantly lower than that of the Responsive group (*t* = -2.898, *p*_Bonferroni_ = 0.013, see Fig.3B), and no significant difference was observed between the HC group and the Responsive group (*t* = -0.302, *p*_Bonferroni_ = 1, see Fig.3B). This result is consistent with the network-level findings. Furthermore, by performing a weighted sum of the individual-level expression matrix, we quantified the integration strength of each ROI. The results showed that the inferior temporal gyrus (ITG), middle frontal gyrus (MFG), orbital gyrus (OrG), parahippocampal gyrus (PhG), and fusiform gyrus (FuG) were key nodes with high integration in the LN-DAN. Similarly, in the ROI-level integration coefficient matrix of the LN-FPN, NMF identified 8 classic coactivation patterns. One-way ANOVA results showed that pattern 6 (pattern#6) exhibited significant differences between the Responsive group, Resistant group, and HC group (*F*_(2, 129)_ = 9.393, *p* < 0.001, see Fig.3C). Further post-hoc two-sample t-tests showed that, after Bonferroni correction, the integration coefficient of the Resistant group was significantly lower than that of the Responsive group (*t* = -4.150, *p*_Bonferroni_ < 0.001, see Fig.3C), and no significant difference was observed between the HC group and the Responsive group (*t* = 0.755, *p*_Bonferroni_ = 1, see Fig.3C). This result is also consistent with the network-level findings. By performing a weighted sum of the individual-level expression matrix, we further quantified the integration strength of each ROI. The results showed that the hippocampus, amygdala, PhG, MFG, and inferior frontal gyrus (IFG) were key nodes with high integration in the LN-FPN. In summary, the NMF-identified coactivation patterns highlight significant alterations in the integration of specific brain regions within the LN, DAN, and FPN that distinguish the Resistant group from the Responsive group and HCs, suggesting a more localized network basis for the observed group differences.

**Figure 3.**
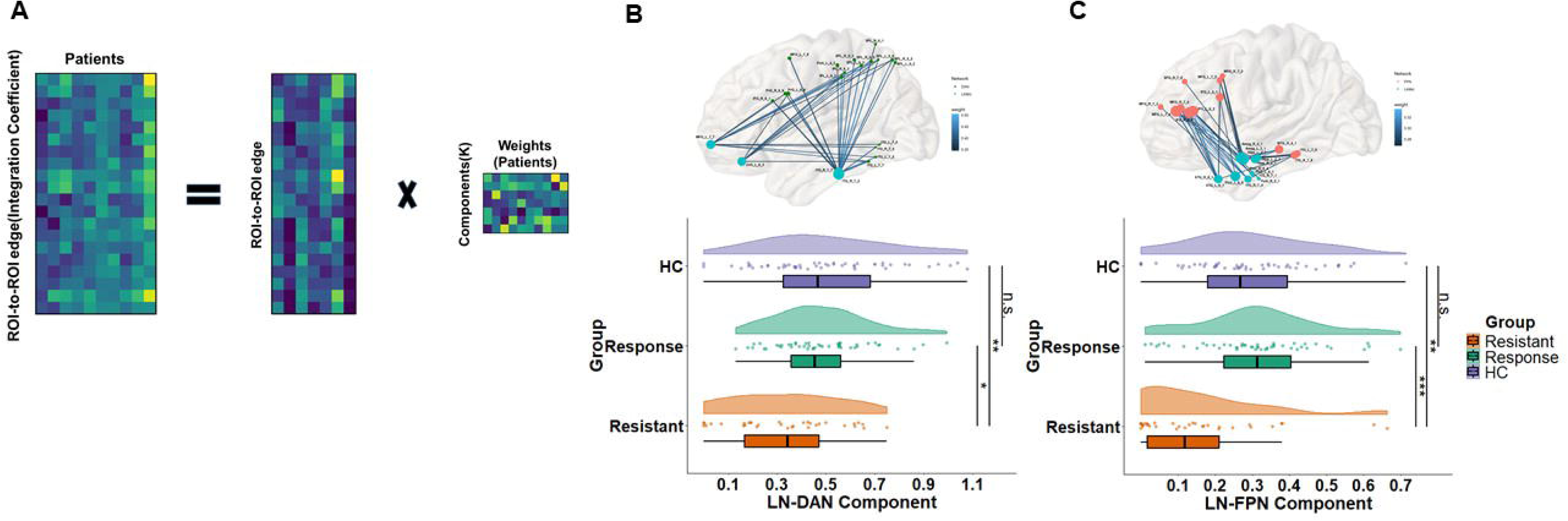
Identification of classic coactivation patterns at the ROI level using NMF (A) Nonnegative matrix factorization (NMF) was applied to the ROI-level integration coefficient matrices of the LN–DAN and LN–FPN (V matrix, with dimensions: patients × ROI-level connections), decomposing each into k group-level integration patterns (W matrix) and corresponding individual-level expression coefficients (H matrix), thereby quantifying the expression of each pattern at the individual level. (B) In the LN–DAN, NMF identified 4 classic coactivation patterns. One-way ANOVA revealed a significant group difference in pattern #1 among HC, the Responsive group, and the Resistant group (*F*_(2,129)_ = 5.823, *p* = 0.004). Post hoc t-tests indicated that the Resistant group showed significantly lower expression of pattern #1 than the Responsive group after Bonferroni correction (*t* = −2.898, *p*_Bonferroni_ = 0.013), while no significant difference was found between HC and the Responsive group (*t* = −0.302, *p*_Bonferroni_ = 1). This result is consistent with network-level findings. The weighted expression analysis identified key high-integration ROIs within this pattern, including the inferior temporal gyrus (ITG), middle frontal gyrus (MFG), orbital gyrus (OrG), parahippocampal gyrus (PhG), and fusiform gyrus (FuG). (C) In the LN–FPN, NMF identified 8 classic coactivation patterns. One-way ANOVA showed a significant group difference in pattern #6 (*F*_(2,129)_ = 9.393, *p* < 0.001). Post hoc comparisons revealed significantly reduced expression in the Resistant group compared to the Responsive group (*t* = −4.150, *p*_Bonferroni_ < 0.001), with no significant difference between HC and the Responsive group (*t* = 0.755, *p*_Bonferroni_ = 1). ROI-level integration strength analysis highlighted the hippocampus, amygdala, PhG, MFG, and inferior frontal gyrus (IFG) as key nodes with high integration within this pattern. \**p* < 0.05, \*\**p* < 0.01, \*\*\**p* < 0.001.

### Correlation and Mediation Analysis

A correlation analysis was conducted between the clinical variable (i.e., SDS score) and the LN-DAN integration coefficient. It was found that in TLE patients, the LN-DAN integration coefficient was significantly negatively correlated with the SDS score (*r* = -0.28, *p* = 0.01, see Fig.4A). Given the significant negative correlation between the LN-DAN integration coefficient and the SDS score, a mediation analysis was performed to examine whether this dynamic reconfiguration could mediate the relationship between group status (Responsive vs. Resistant) and SDS. The results indicated that the indirect path through the lower LN-DAN integration coefficient (a*b, beta = 1.647, standard error = 0.045; 95% confidence interval = [0.005, 0.179], *Z* = 36.751, *p* < 0.001, mediation = 35.81%, see Fig.4B) fully mediated the direct path from group status to SDS (c’, beta = 2.952, standard error = 2.195; 95% confidence interval = [-1.350, 7.255], *Z* = 1.345, *p* = 0.183, mediation = 64.19%, see Fig.4B). That is, compared to the Responsive group, the Resistant group exhibited lower DAN-LN integration coefficients, which in turn was associated with higher levels of depression.

**Figure 4.**
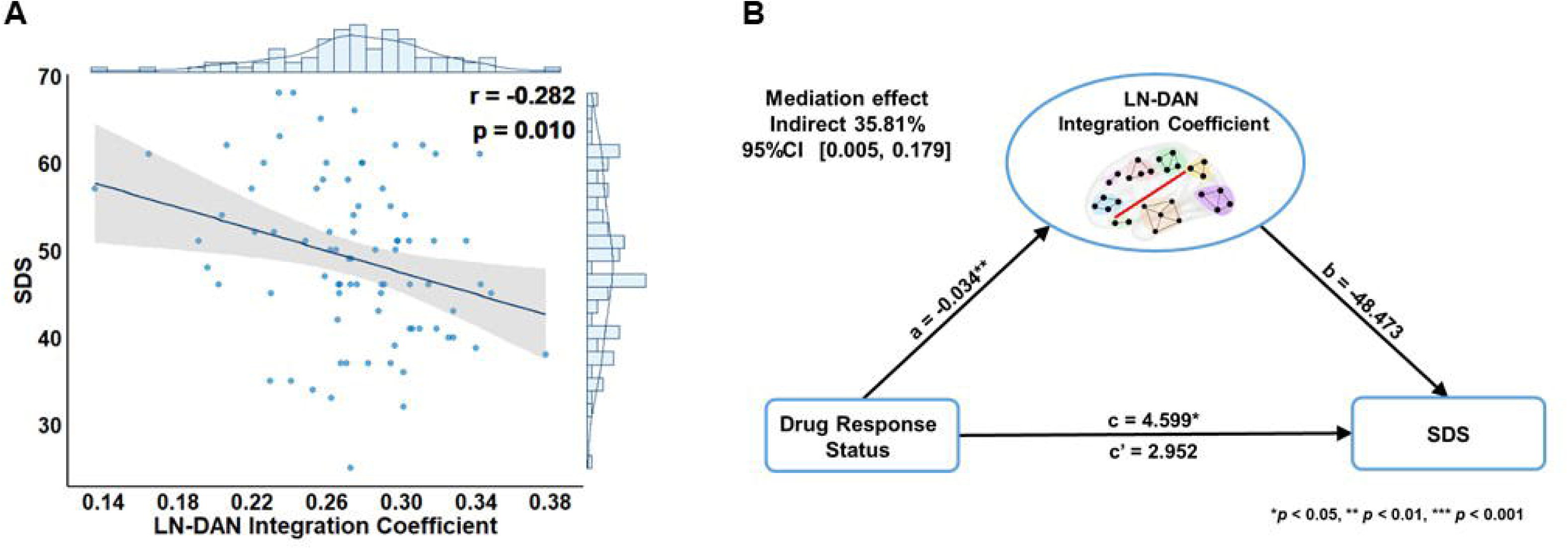
Mediating role of LN–DAN integration in the relationship between Drug Response Status and depressive symptoms (A) Among TLE patients, the LN–DAN integration coefficient was significantly negatively correlated with SDS scores (*r* = −0.282, *p* = 0.010), suggesting that lower network integration is associated with more severe depressive symptoms. (B) Given the observed correlation, a mediation analysis was conducted to examine whether LN–DAN integration mediates the relationship between group status (Resistant vs. Responsive) and SDS scores. The results revealed a significant full mediation effect: the indirect pathway through LN–DAN integration was significant (a × b, *β* = 1.647, standard error = 0.045, 95% CI = [0.005, 0.179], *Z* = 36.751, *p* < 0.001), accounting for 35.81% of the total effect. The direct pathway (c′) was not significant (*β* = 2.952, standard error = 2.195, 95% CI = [−1.350, 7.255], *Z* = 1.345, *p* = 0.183), indicating that LN–DAN integration plays a mediating role in the link between group status and depressive symptoms. \**p* < 0.05, \*\**p* < 0.01, \*\*\**p* < 0.001.

### Neurotransmitter Receptor Correlates of Drug Response in TLE

Spatial correlation analysis between neurotransmitter receptor distribution and neuroimaging revealed that the difference in the integration coefficients of the LN (LN-DAN, LN-FPN) between the Responsive group and Resistant group was positively correlated with the density of 5-HT1B receptors (*r* = 0.38, *p* = 0.003, *q* = 0.038, see Fig.5B) and negatively correlated with the density of D2 receptors (*r* = -0.43, *p* = 0.004, *q* = 0.038, see Fig.5B). This finding may suggest that 5-HT1B and D2 receptors play a key role in the neurobiological mechanisms that differentiate the Resistant group from the Responsive group.

**Figure 5.**
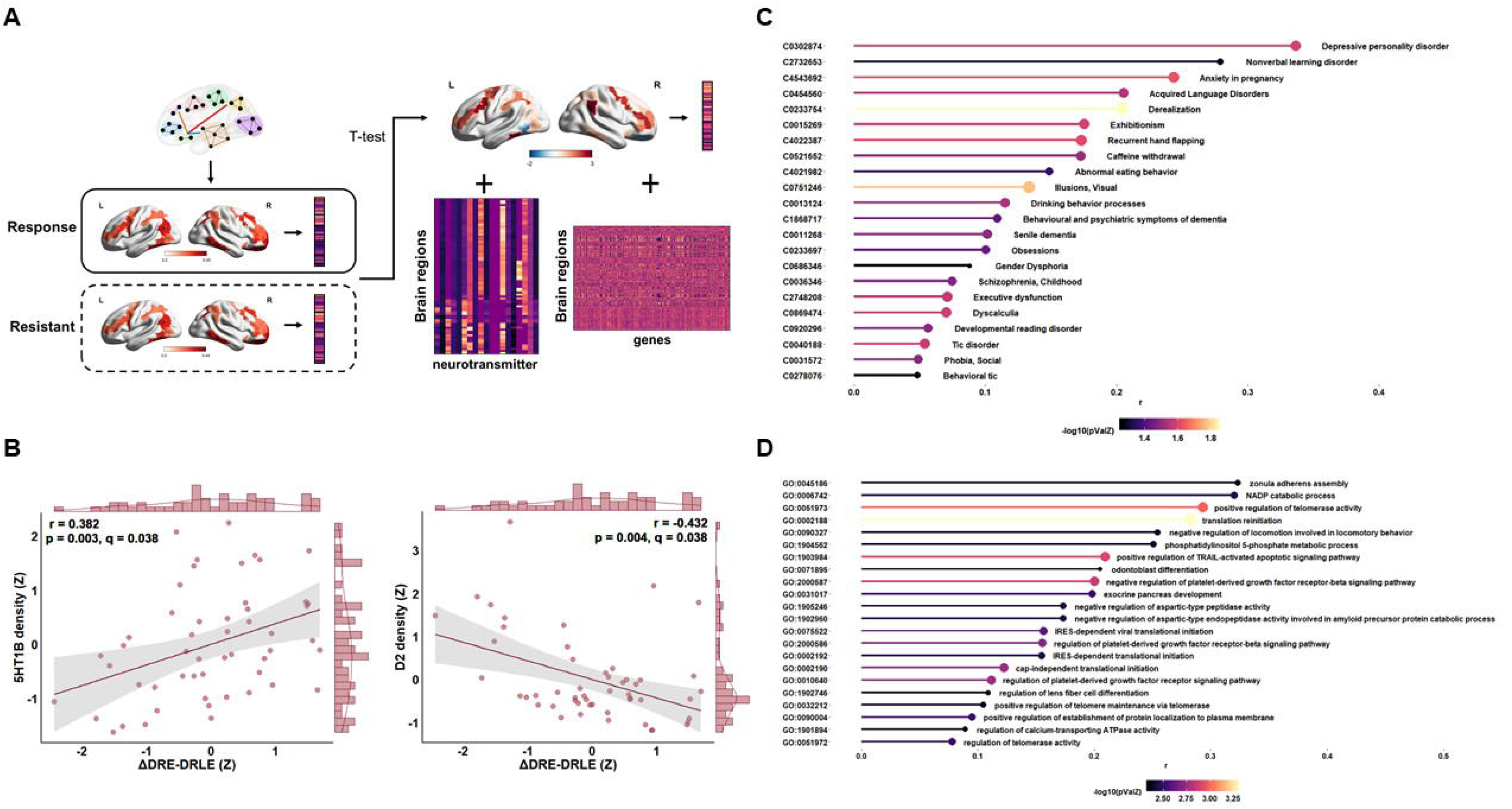
Spatial associations between LN integration differences and neurotransmitter receptor density, genetic disease markers, and functional gene pathways (A) The integration strength between the LN and 52 ROIs within the DAN and FPN networks was compared between the Responsive group and Resistant group using independent-sample t-tests. The resulting t-statistics were then used for spatial correlation analyses with neurotransmitter/transporter distribution maps and gene expression profiles. (B) Spatial correlation analysis between neurotransmitter receptor distribution and neuroimaging data revealed that differences in LN integration coefficients (including LN–DAN and LN–FPN) between the Responsive group and Resistant group were positively correlated with the density of 5-HT1B receptors (*r* = 0.38, *p* = 0.003, *q* = 0.038) and negatively correlated with the density of D2 receptors (*r* = −0.43, *p* = 0.004, *q* = 0.038). (C) Spatial correlation analysis between neuroimaging findings and genetic disease risk markers showed that LN integration differences were strongly associated with genes implicated in affective disorders (e.g., Depressive personality disorder: *r* = 0.337, *p* = 0.021; Nonverbal learning disorder: *r* = 0.279, *p* = 0.048; Anxiety in pregnancy: *r* = 0.244, *p* = 0.022). These results were significant before multiple comparison correction. (D) Further analyses revealed that LN integration differences were significantly positively correlated with several functional gene pathways, including positive regulation of telomerase activity (cScorePheno = 0.293, *p* < 0.001, *q* < 0.001), translation reinitiation (cScorePheno = 0.283, *p* < 0.001, *q* < 0.001), positive regulation of TRAIL-activated apoptotic signaling pathway (cScorePheno = 0.210, *p* < 0.001, *q* < 0.001), and cap-independent translational initiation (cScorePheno = 0.123, *p* < 0.001, *q* < 0.001).

### Spatial Associations with Genetic Disease Markers and Molecular Processes

Spatial correlation analysis between genetic disease markers and neuroimaging revealed that the difference in the integration coefficients of the LN between the Responsive group and Resistant group was closely related to genes associated with depression (e.g., Depressive personality disorder: *r* = 0.337, *p* = 0.021; Nonverbal learning disorder: *r* = 0.279, *p* = 0.048; Anxiety pregnancy: *r* = 0.244, *p* = 0.022, see Fig.5C). Notably, the difference in the LN integration coefficients between the Responsive group and Resistant group was significantly positively correlated with the Positive regulation of telomerase activity (cScorePheno = 0.293, *p* < 0.001, *q* < 0.001, see Fig.5C), Translation reinitiation (cScorePheno = 0.283, *p* < 0.001, *q* < 0.001, see Fig.5C), Positive regulation of TRAIL-activated apoptotic signaling pathway (cScorePheno = 0.210, *p* < 0.001, *q* < 0.001, see Fig.5C), and Cap-independent translational initiation (cScorePheno = 0.123, *p* < 0.001, *q* < 0.001, see Fig.5C). These results begin to delineate molecular mechanisms that differ between the Responsive group and Resistant group, offering insights into the biological connections that may underlie the observed links between drug resistance, altered brain function, and comorbid depression in epilepsy.

## Discussion

By integrating dynamic brain network analysis, NMF, neuroimaging-transcriptomics, and neurotransmitter spatial mapping, our study provides robust multi-level evidence suggesting that drug resistance in TLE is linked to a marked exacerbation of comorbid depressive symptoms. Crucially, this more severe presentation of depression appears to be associated with specific alterations in the dynamic integration between the LN and the DAN, along with distinct neurochemical and transcriptomic signatures, highlighting a systems-level mechanism that links aberrant network dynamics to affective comorbidity. The results showed that, compared to the Responsive group, patients in the Resistant group exhibited more pronounced functional abnormalities in the LN. Significant differences were found in the integration coefficients between the LN and the DAN and the FPN, and crucially, the DAN-LN integration degree mediated the relationship between group status and depressive symptoms. Further coactivation pattern analysis identified key coactivation patterns and nodes (such as the hippocampus, amygdala, and MFG) in the LN-DAN and LN-FPN, with significant differences between the Responsive and Resistant groups, indicative of more severe network disruption in the resistant state. Moreover, we found that the density of 5-HT1B and D2 receptors was significantly correlated with differences in LN integration between the Responsive and Resistant groups, suggesting that these neurotransmitter systems may be involved in the neurobiological mechanisms through which drug response affects brain function and emotional regulation. Concurrently, the differences in LN functional integration were significantly associated with the expression levels of various depression-related genes and cellular signaling pathways (such as telomerase activity regulation, TRAIL-mediated apoptosis signaling pathway, etc.). This study therefore not only emphasizes the critical role of the LN in DRE and its comorbid depression but, by for the first time linking these dynamic functional changes to specific neurochemical and molecular substrates in the context of drug resistance, also provides insights into a potential indirect mechanism through which drug response status is associated with depressive psychopathology, suggesting new potential targets for future precision therapies aimed at mitigating this amplified burden.

### Altered LN Dynamic Integration Associated with Drug Response

The LN is pivotal for emotional regulation, memory, and cognition, and its dysfunction is prominent in DRE patients, often accompanying emotional and cognitive deficits.^40,41^ Our analysis revealed significant differences in the dynamic integration coefficients of the LN (interacting with the DAN and FPN) among HC, the Responsive group, and the Resistant group, underscoring the centrality of LN dynamic integration in the mechanisms differentiating drug response status in these TLE patients. Specifically, patients in the Resistant group exhibited significantly lower LN integration compared to patients in the Responsive group. This may reflect the consequence of long-term, excessive neuronal synchrony impacting LN stability, potentially associated with imbalances in emotional and cognitive regulation that likely relate to amplified depressive symptoms. The reduced LN-DAN integration in the Resistant group suggests that in the drug-resistant state, the LN’s capacity for emotional regulation might be compromised by its interaction with the DAN. This impaired dynamic balance between networks crucial for attention and emotion appears closely related to the worsening of depressive symptoms, highlighting the importance of DAN-LN interactions in maintaining emotional stability.^42–43^ Similarly, decreased LN-FPN integration in patients in the Resistant group points to an imbalance between cognitive control and emotional processing, which can further exacerbate mood disturbances. The Resistant group’s distinct profile across these networks, all vital for attention, emotion, and cognition, suggests that their drug-resistant state is associated with more severe and widespread dynamic network dysregulation, which in turn is linked to a greater burden of depression.

In contrast, the LN integration profile of the Responsive group more closely resembled that of HCs. This suggests that in the context of an effective response to antiseizure medication, as seen in the Responsive group, these dynamic network disruptions appear to be less severe or partially mitigated, potentially fostering recovery of emotional and cognitive functions. These observations underscore the potential of dynamic network integration as a measure of treatment effectiveness and suggest that interventions for DRE with comorbid depression might benefit from targeting cross-network functional reorganization.

### Key Coactivation Patterns and Nodal Contributions Associated with Drug Response

To pinpoint specific regional contributions to these network-level findings, we employed NMF. This approach identified several coactivation patterns within the LN-DAN and LN-FPN, among which certain patterns showed significant differences between the Responsive and Resistant groups, consistent with the alterations observed at the network level. Within a key LN-DAN coactivation pattern, nodes such as the ITG, MFG, OrG, PhG, and FuG showed strong integration. These regions are integral to emotional processing, attention, and memory; our findings support their role in cognitive dysfunction in epilepsy^44^ and suggest their abnormal integration patterns may contribute to drug treatment insensitivity in DRE. Similarly, within the LN-FPN interactions, a coactivation pattern involving high integration of the hippocampus, amygdala, PhG, MFG, and IFG was significantly different in patients in the Resistant group. The hippocampus and amygdala are well-recognized for their sensitivity and central role in epileptic seizures and drug resistance.^45^ Our results suggest that the drug-resistant state further compromises the dynamic functional integration of these critical emotional and memory hubs, potentially contributing to the more severe depressive phenotype observed in patients in the Resistant group.

The consistency between ROI-level NMF findings and network-level analyses validates the importance of dynamic network functional integration in mechanisms differentiating drug response states.

### DAN-LN Integration Mediates the Link Between Drug Response Status and Depression

This study found that the integration of the DAN-LN significantly mediated the relationship between group status and depressive symptoms. This result suggests that the link between a favorable drug response and milder depressive symptoms is not a direct, but is significantly associated with the dynamic reconfiguration of the DAN-LN. In the Responsive group, the functional reconfiguration of the DAN-LN may be associated with a restoration of the dynamic interaction between the DAN and the LN, which could potentially relate to an alleviation of depressive symptoms. This finding is consistent with previous research in depression, where evidence has shown that patients with depression exhibit DAN dysfunction^46^ and abnormal dynamic interactions across multiple brain networks.^21^ Furthermore, in studies of DRE, Xie et al. found that in patients with drug-resistant focal cortical dysplasia-related epilepsy (FCD-PRE), the functional connectivity of the DAN was significantly reduced, while enhanced connectivity between the DAN and the LN was associated with better treatment outcomes.^47^ This finding supports the importance of the DAN-LN in differentiating drug response outcomes. Our results therefore suggest that DAN-LN integration may serve as a neurobiological marker reflecting the neurophysiological differences associated with drug response status, with its disruption in patients in the Resistant group representing a key factor associated with more severe depressive symptoms.

### Neurochemical and Molecular Correlates of Altered LN Dynamics

We found that differences in LN integration coefficients between Responsive and Resistant groups were significantly correlated with the spatial density of serotonin 5-HT1B and dopamine D2 receptors. The 5-HT1B receptor, an inhibitory serotonin receptor highly expressed in limbic and frontal regions, influences epilepsy by regulating neuronal excitability and network stability.^48, 49^ Its role in depression and antidepressant response is also established, with hippocampal 5-HT1B binding levels correlating with treatment efficacy.^50^ Our findings suggest that reduced 5-HT1B receptor density in critical brain areas might be linked to decreased inhibitory control over the LN in DRE, potentially relating to compromised network stability and an association with more severe depressive symptoms alongside poor drug response. Similarly, D2 receptors, integral to dopamine signaling within the LN, are implicated in emotional and cognitive regulation, and their dysfunction is linked to epilepsy and depression.^51, 52^ Higher D2 receptor density in the Resistant group might reflect dysregulated dopamine signaling within the LN, which could be further associated with impaired functional integration and linked to affective disturbances. These findings implicate 5-HT1B and D2 receptor systems in modulating LN dynamic stability associated with drug response, offering potential targets for future therapies.

Furthermore, spatial association analyses linked LN integration differences to the expression of depression-related genes and molecular pathways, including those involved in telomerase activity regulation and apoptosis signaling (TRAIL-activated pathway). Genes associated with phenotypes such as depressive personality disorder showed correlations with LN integration differences, suggesting a genetic influence on the functional LN reconfiguration observed in our TLE patients, which in turn may affect clinical depressive presentations. The LN is central to emotional regulation and its function is modulated by neurotransmitter signaling and neuroplasticity.^53,54^ Our study suggest that in TLE, specific gene expression patterns show associations with depressive symptom severity and LN functional homeostasis, which in turn relate to individual differences in drug responsiveness. The significant correlation with telomerase activity, with its known role in depression and neurodegeneration,^55, 56^ and TRAIL signaling, essential for neuronal apoptosis (Henshall, 2007), strongly suggests these molecular processes are integral to the pathophysiology linking LN dynamic changes, drug response status, and the exacerbation of depressive symptoms in TLE.

### Limitations

It is important to note that our study has several limitations. First, the cross-sectional design of this study precludes causal inferences and the tracking of dynamic changes in relation to treatment initiation or the progressive development of drug resistance and mood changes. Longitudinal studies are needed to clarify these temporal dynamics. Second, while we controlled for age and gender, other potential confounders, such as illness duration, specific antiseizure medication history, and other comorbidities, might have influenced the observed differences in LN integration; future studies should aim for more comprehensive control of such factors. Third, the neurotransmitter receptor density maps and our neuroimaging data were derived from different samples, and the receptor maps had a relatively small sample size with uneven gender distribution, potentially introducing bias and limiting generalizability. Fourth, the correlational nature of our analyses, including those with receptor densities and gene expression, does not establish causality; interventional studies are required to explore the causal roles of these neurotransmitter and molecular systems. Finally, individual heterogeneity in gene expression and its spatial correlation with imaging findings can lead to variability; larger, gender-balanced samples are desirable to enhance the robustness of these associations.

## Conclusions

This study found that the LN and its integration coefficients with the DAN and the FPN could serve as effective neural biomarkers to distinguish drug response status in TLE patients. The dynamic reconfiguration of the LN is closely related to neurotransmitter receptor regulation, gene expression, and molecular processes, suggesting that drug response status in epilepsy may be associated with plastic changes in neural networks and functional reorganization of the neurotransmitter systems. These synaptic and molecular-level changes, particularly those changes reflecting drug response status, may constitute the potential neuropathological basis for the often exacerbated emotional disorders, notably depression, observed in TLE patients with drug resistance. Understanding these alterations is crucial for developing strategies to mitigate the compounded burden of drug resistance and comorbid depression.

## Data availability statement

The data that support the findings of this study are available from the corresponding author upon reasonable request.

## Author Contribution

Z.W., H.T., X.Y., and X.H. contributed to the conception and design of the study; Z.W., H.T., L.H., M.W., Y.G., A.C., Y.G., J.H., X.Y., and X.H. contributed to the acquisition and analysis of data; M.W., Y.G., A.C., Y.G., J.H., X.Y., and X.H. contributed to drafting the text or preparing the figures.

## REFERENCES

1. Kanner AM. Is depression associated with an increased risk of treatment-resistant epilepsy? Research strategies to investigate this question. Epilepsy Behav 2014;38:3–7.

2. Jansen C, Francomme L, Vignal J-P, et al. Interictal psychiatric comorbidities of drug-resistant focal epilepsy: prevalence and influence of the localization of the epilepsy. Epilepsy & behavior 2019;94:288–296.

3. Kanner AM. Management of psychiatric and neurological comorbidities in epilepsy. Nature Reviews Neurology 2016;12:106–116.

4. Zheng Y, Ding X, Guo Y, et al. Multidisciplinary management improves anxiety, depression, medication adherence, and quality of life among patients with epilepsy in eastern China: A prospective study. Epilepsy & Behavior 2019;100:106400.

5. Błaszczyk B, Czuczwar SJ. Epilepsy coexisting with depression. Pharmacological reports 2016;68:1084–1092.

6. Kanner AM, Ribot R, Mazarati A. Bidirectional relations among common psychiatric and neurologic comorbidities and epilepsy: Do they have an impact on the course of the seizure disorder? Epilepsia open 2018;3:210–219.

7. Kanner AM. Depression and epilepsy: a review of multiple facets of their close relation. Neurologic clinics 2009;27:865–880.

8. Chen S, Wu X, Lui S, et al. Resting-state fMRI study of treatment-naive temporal lobe epilepsy patients with depressive symptoms. Neuroimage 2012;60:299–304.

9. Kanner AM, Schachter SC, Barry JJ, et al. Depression and epilepsy: epidemiologic and neurobiologic perspectives that may explain their high comorbid occurrence. Epilepsy & Behavior 2012;24:156–168.

10. Hecimovic H, Santos J, Price JL, et al. Severe hippocampal atrophy is not associated with depression in temporal lobe epilepsy. Epilepsy & behavior 2014;34:9–14.

11. Elkommos S, Mula M. A systematic review of neuroimaging studies of depression in adults with epilepsy. Epilepsy & Behavior 2021;115:107695.

12. Peng W, Mao L, Yin D, et al. Functional network changes in the hippocampus contribute to depressive symptoms in epilepsy. Seizure 2018;60:16–22.

13. Josephson CB, Lowerison M, Vallerand I, et al. Association of depression and treated depression with epilepsy and seizure outcomes: a multicohort analysis. JAMA neurology 2017;74:533–539.

14. Scott AJ, Sharpe L, Hunt C, Gandy M. Anxiety and depressive disorders in people with epilepsy: a meta-analysis. Epilepsia 2017;58:973–982.

15. Kalilani L, Sun X, Pelgrims B, et al. The epidemiology of drug-resistant epilepsy: a systematic review and meta-analysis. Epilepsia 2018;59:2179–2193.

16. Tang F, Hartz AM, Bauer B. Drug-resistant epilepsy: multiple hypotheses, few answers. Frontiers in neurology 2017;8:301.

17. Perucca E, Perucca P, White HS, Wirrell EC. Drug resistance in epilepsy. Lancet Neurol 2023;22:723–734.

18. Beghi E, Beretta S, Carone D, et al. Prognostic patterns and predictors in epilepsy: a multicentre study (PRO-LONG). Journal of Neurology, Neurosurgery & Psychiatry 2019;90:1276–1285.

19. Demirtaş M, Tornador C, Falcón C, et al. Dynamic functional connectivity reveals altered variability in functional connectivity among patients with major depressive disorder. Human brain mapping 2016;37:2918–2930.

20. Shunkai L, Su T, Zhong S, et al. Abnormal dynamic functional connectivity of hippocampal subregions associated with working memory impairment in melancholic depression. Psychological medicine 2023;53:2923–2935.

21. Tian S, Chattun MR, Zhang S, et al. Dynamic community structure in major depressive disorder: A resting-state MEG study. Progress in Neuro-Psychopharmacology and Biological Psychiatry 2019;92:39–47.

22. Fisher RS, van Emde Boas W, Blume W, et al. Epileptische Anfälle und Epilepsie: von der Internationalen Liga gegen Epilepsie (International League Against Epilepsy; ILAE) und dem Internationalen Büro für Epilepsie (International Bureau for Epilepsy; IBE) vorgeschlagene Definitionen. Aktuelle Neurologie 2005;32:249–252.

23. Zung WW, Richards CB, Short MJ. Self-rating depression scale in an outpatient clinic: further validation of the SDS. Archives of general psychiatry 1965;13:508–515.

24. Kwan P, Arzimanoglou A, Berg AT, et al. Definition of drug resistant epilepsy: consensus proposal by the ad hoc Task Force of the ILAE Commission on Therapeutic Strategies. Wiley Online Library; 2010.

25. Esteban O, Markiewicz CJ, Blair RW, et al. fMRIPrep: a robust preprocessing pipeline for functional MRI. Nature methods 2019;16:111–116.

26. Mehta K, Salo T, Madison TJ, et al. XCP-D: A Robust Pipeline for the post-processing of fMRI data. Imaging Neuroscience 2024;2:1–26.

27. Wang J, Wang X, Xia M, et al. GRETNA: a graph theoretical network analysis toolbox for imaging connectomics. Frontiers in human neuroscience 2015;9:386.

28. Fan L, Li H, Zhuo J, et al. The human brainnetome atlas: a new brain atlas based on connectional architecture. Cerebral cortex 2016;26:3508–3526.

29. Newman ME. Modularity and community structure in networks. Proceedings of the national academy of sciences 2006;103:8577–8582.

30. He X, Bassett DS, Chaitanya G, et al. Disrupted dynamic network reconfiguration of the language system in temporal lobe epilepsy. Brain 2018;141:1375–1389.

31. Hayes AF. PROCESS: A versatile computational tool for observed variable mediation, moderation, and conditional process modeling. University of Kansas, KS; 2012.

32. Lee DD, Seung HS. Learning the parts of objects by non-negative matrix factorization. Nature 1999;401:788–791.

33. Zhou T, Kang J, Cong F, Li X. Stability-driven non-negative matrix factorization-based approach for extracting dynamic network from resting-state EEG. Neurocomputing 2020;389:123–131.

34. Li H, Satterthwaite TD, Fan Y. Large-scale sparse functional networks from resting state fMRI. Neuroimage 2017;156:1–13.

35. Hansen JY, Shafiei G, Markello RD, et al. Mapping neurotransmitter systems to the structural and functional organization of the human neocortex. Nature neuroscience 2022;25:1569–1581.

36. Hawrylycz MJ, Lein ES, Guillozet-Bongaarts AL, et al. An anatomically comprehensive atlas of the adult human brain transcriptome. Nature 2012;489:391–399.

37. Arnatkevic̆iūtė A, Fulcher BD, Fornito A. A practical guide to linking brain-wide gene expression and neuroimaging data. Neuroimage 2019;189:353–367.

38. Habets PC, Kalafatakis K, Dzyubachyk O, et al. Transcriptional and cell type profiles of cortical brain regions showing ultradian cortisol rhythm dependent responses to emotional face stimulation. Neurobiology of Stress 2023;22:100514.

39. Lotter L, Dukart J, Fulcher B. ABAnnotate: A toolbox for ensemble-based multimodal gene-category enrichment analysis of human neuroimaging data. Published online April 2022;15.

40. Wang K, Xie F, Liu C, et al. Abnormal functional connectivity profiles predict drug responsiveness in patients with temporal lobe epilepsy. Epilepsia 2022;63:463–473.

41. Aung T, Tang LW, Ho J, et al. Differential functional connectivity of amygdala in drug-resistant temporal lobe epilepsy. Epilepsia 2025

42. 41.Viviani R. Emotion regulation, attention to emotion, and the ventral attentional network. Frontiers in human neuroscience 2013;7:746.

43. Kohn N, Eickhoff SB, Scheller M, et al. Neural network of cognitive emotion regulation—an ALE meta-analysis and MACM analysis. Neuroimage 2014;87:345–355.

44. Wang H, Tan G, Li X, et al. Aberrant functional connectivity associated with drug response in patients with newly diagnosed epilepsy. Neurological Sciences 2024;45:4973–4982.

45. Pitkänen A, Löscher W, Vezzani A, et al. Advances in the development of biomarkers for epilepsy. The Lancet Neurology 2016;15:843–856.

46. Yang H, Chen X, Chen Z-B, et al. Disrupted intrinsic functional brain topology in patients with major depressive disorder. Molecular psychiatry 2021;26:7363–7371.

47. Xie H, Illapani VSP, Vezina LG, et al. Mapping Functional Connectivity Signatures of Pharmacoresistant Focal Cortical Dysplasia-Related Epilepsy. Annals of neurology 2025;97:51–61.

48. Wesołowska A, Nikiforuk A, Chojnacka-Wójcik E. Anticonvulsant effect of the selective 5-HT1B receptor agonist CP 94253 in mice. European journal of pharmacology 2006;541:57–63.

49. Jaworski T. Control of neuronal excitability by GSK-3beta: Epilepsy and beyond. Biochimica et Biophysica Acta (BBA)-Molecular Cell Research 2020;1867:118745.

50. Tiger M, Veldman ER, Ekman C-J, et al. A randomized placebo-controlled PET study of ketaminés effect on serotonin1B receptor binding in patients with SSRI-resistant depression. Translational psychiatry 2020;10:159.

51. Zarcone D, Corbetta S. Shared mechanisms of epilepsy, migraine and affective disorders. Neurological Sciences 2017;38:73–76.

52. Rezaei M, Sadeghian A, Roohi N, et al. Epilepsy and dopaminergic system. Physiology and Pharmacology 2017;21:1–14.

53. Tartt AN, Mariani MB, Hen R, et al. Dysregulation of adult hippocampal neuroplasticity in major depression: pathogenesis and therapeutic implications. Molecular psychiatry 2022;27:2689–2699.

54. Jiang Y, Zou D, Li Y, et al. Monoamine neurotransmitters control basic emotions and affect major depressive disorders. Pharmaceuticals 2022;15:1203.

55. Zhou Q-G, Hu Y, Wu D-L, et al. Hippocampal telomerase is involved in the modulation of depressive behaviors. Journal of Neuroscience 2011;31:12258–12269.

56. Wang J, Liu Y, Xia Q, et al. Potential roles of telomeres and telomerase in neurodegenerative diseases. International journal of biological macromolecules 2020;163:1060–1078.

